# HIV-1 subtype C LTR Sp1IIIT5A mutant enhances transcription activity and Sp1 binding affinity

**DOI:** 10.1101/2025.01.23.634640

**Authors:** Nomcebo Msthali, Ibrahim Kehinde, Rene Khan, Mahmoud Soliman, Thumbi Ndung’u, Paradise Madlala

**Affiliations:** HIV Pathogenesis Programme, The Doris Duke Medical Research Institute, Nelson R. Mandela School of Medicine, University of KwaZulu-Natal, Durban, South Africa; School of Laboratory Medicine and Medical Sciences, University of KwaZulu-Natal, Durban, South Africa; Molecular Bio-Computation and Drug Design Lab, School of Health Sciences, University of KwaZulu-Natal, Durban 4000, South Africa; Discipline of Medical Biochemistry, School of Laboratory Medicine and Medical Science, College of Health Sciences, University of KwaZulu-Natal, Durban, South Africa; Ragon Institute of Massachusetts General Hospital, Massachusetts Institute of Technology, and Harvard University, Cambridge, Massachusetts, United States of America; Africa Health Research Institute (AHRI), Durban, South Africa; Division of Infection and Immunity, University College London, London, United Kingdom

## Abstract

**Background:** Genetic variation within HIV-1 subtype C (HIV-1C) long terminal repeat (LTR) transmitted/founder viruses influences transcription activation and clinical disease outcomes. The role of specific mutations such as thymine-to-adenine (T5A) mutation at position five of the Specificity protein 1 (Sp1) III motif (Sp1IIIT5A) remains underexplored. This study investigates the impact of Sp1IIIT5A on HIV-1C LTR transcription activity and Sp1 binding affinity.

**Methods:** The Sp1IIIT5A mutant and consensus HIV-1C LTR sequences were cloned into the pGL3 Luciferase Basic reporter vector, sequenced, and transfected into SVG and Jurkat cell lines, independently. Transcription activity and Sp1 expression were assessed via luciferase assays and Western blot. Structural models of Sp1IIIT5A, consensus LTRs and Sp1 were generated, and docking scores calculated using HDOCK, HADDOCK, and pyDockDNA. Molecular dynamics simulations analyzed stability and interactions of Sp1IIIT5A LTR-Sp1 complexes.

**Results and Discussion:** The Sp1III5A mutant significantly increased basal (SVG: p<0.0001; Jurkat: p=0.0052) and Tat-mediated (SVG and Jurkat: p<0.0001) HIV-1C LTR transcription activity in both cell lines, with stronger effects in SVG cells. Sp1 expression levels remained similar across cell lines (p=0.0814). Sp1III5A exhibited higher binding affinity (-332.7, -174.6, and -279.2 kcal/mol) than the canonical sequence (-311.4, -157.0, and -247.3 kcal/mol).

**Conclusion:** The Sp1IIIT5A mutation significantly enhances HIV-1C LTR transcription activity and Sp1 binding affinity, indicating its potential tole in modulating HIV-1C transcription and pathogenesis. Further investigation is needed to elucidate its impact on HIV-1C latency.

**Importance:** In this study we show that the thymine-to-adenine (T5A) mutation at position five of the Sp1 III motif (Sp1IIIT5A) within the HIV-1 subtype C (HIV-1C) long terminal repeat (LTR) increases viral transcription. This mutation enhances the interaction between HIV-1C and the cellular transcription factor Sp1, promoting the viral strain’s ability to replicate. Our findings provide insight into why certain HIV-1C strains behave differently, potentially leading to heterogenous rates of disease progression. Understanding the Sp1IIIT5A mutation could lead to improved strategies for controlling HIV-1C and developing cure strategies to clear the infection or result in virus remission.

## Introduction

People living with HIV (PLWH) who are antiretroviral therapy naïve have widely differing rates of clinical disease progression (1). The underlying mechanisms for this heterogeneity includes host and viral factors but these are not completely understood. The viral promoter, 5’ long terminal repeat (LTR) drives gene transcription and is essential for the viral life cycle (2, 3). HIV-1 5’ LTR is divided into three regions: Unique 3 (U3), Repeat (R) and Unique 5 (U5) regions, with the U3 region further subdivided into modulatory, core-enhancer and core-promoter domains (reviewed in (4)). Each of these domains contains distinct transcription factor binding sites (TFBSs); notably, the core-promoter domain features the TATA box, two E-boxes, and three Specificity protein 1 (Sp1) binding sites, denoted as Sp1I, Sp1II and Sp1III (reviewed in (4)). Sp1 is a transcription factor family, which participates in the regulation of cellular gene expression and is required to initiate HIV-1 transcription by binding to the Sp1 motifs located within the 5’ LTR (5). Subsequently, Sp1 interacts with TATA binding protein (TBP) and some TBP associated factors such as TAF110, which are components of the general RNA Polymerase II (RNAPII) transcriptional factor D (6, 7).

Therefore, Sp1 is an essential transcription factors that initiates basal transcription by recruiting the RNAPII-dependent transcriptional machinery to the HIV-1 promoter 5’ LTR (8–10). Short abortive viral mRNA transcripts are produced during basal transcription when the viral protein, Trans-activator of transcription (Tat), which enhances the ability of HIV-1 5’ LTR to drive the expression of full-length HIV-1 proviral transcripts, is absent (11, 12). Taken together, these studies suggest that HIV-1 LTR transcription activity and viral replication are partly dependent on interactions of the Sp1 motifs with the family of transcription factors.

Immune pressures within the host and the error-prone Reverse Transcriptase cause genetic variation within the HIV-1 genome, including the LTR, resulting in viral quasispecies that can be differentially regulated (reviewed in (13)). Inter- and intra-subtype LTR genetic variation translates to functional differences (14, 15). Previous studies reported that single point mutations in the core-promoter region, including Sp1 and Tata Box motifs, dysregulated viral gene expression (16, 17). One of the studies further reported that Sp1 mutations impaired both basal, in the absence of Tat, and Tat-mediated LTR transcription activity (16). A different study identified single nucleotide polymorphisms (SNPs), cytosine-to-thymine (C3T) at position 3 of the C/EBP motif and cytosine-to-thymine (C5T) at position 5 of Sp1III motif that altered 5’ LTR transcription activity (18). Both C/EBP3T and Sp1III5T LTR mutants were able to drive gene expression albeit at low level, suggesting that these mutations could result in reduced binding by their respective cellular transcription factors (18).

Another study reported that the Sp1IIIC5T SNP increased while Sp1IIC5T SNP decreased in frequency during the course of HIV-1 disease, suggesting that LTR Sp1 binding site sequence variants may predict disease outcome (19). All the aforementioned research, however, focused on HIV-1 subtype B, which is accounts for only 12% of all HIV-1 infections worldwide and is most common in Europe and the US. Therefore, a comparable study is required for HIV-1 subtype C (HIV-1C), which is responsible for approximately 50% of the global and 98% of southern Africa infections (20, 21).

Recently, our group showed that genetic variation of the HIV-1C transmitted/founder (T/F) LTR impacts transcription activation and clinical disease outcomes in PLWH from South Africa (22). While most motifs in the T/F LTRs were relatively conserved, the Sp1III motif exhibited high genetic variation where G2A and T5A SNPs were more prevalent, with T5A occurring in 61% of the T/F LTRs that were analyzed (22). However, it remains unclear whether this differential LTR transcription activity and disease outcome is due to a single SNP within the Sp1III motif or a combination of mutations. Therefore, we hypothesize that the single mutation Sp1 IIIT5A in HIV-1C LTR impacts transcription activity through altered Sp1 binding affinity. In this study we demonstrate that the Sp1IIIT5A mutation in HIV-1C LTR significantly enhances transcription activity and binding affinity of Sp1, in both SVG and Jurkat cell lines. These results highlight the potential and crucial role of Sp1IIIT5A in modulating HIV-1 transcription, emphasizing the need for further investigation into its implications for HIV-1C latency.

## RESULTS

### Introduction of Sp1III5A mutation into the HIV-1C subtype C LTR

The previous study conducted by our group demonstrated that certain elements, namely RBE III, TATA Box, NF-кB I, NF-кB II, Sp1I, Sp1II, and the E-Box TFBS, remained relatively unchanged within the LTR regions (22). Consistent with previous reports (18, 19), our group observed noticeable variation within the Sp1III motif displaying high frequences of G2A and T5A SNPs in the T/F LTR variants (22). Specifically, we showed that Sp1IIIT5A SNP occurred in 61% of the T/F LTR variants that were analyzed. Other previous studies had reported that the SNP Sp1IIIC5T among HIV-1 subtype B viruses altered LTR-driven gene transcription (18, 19) and correlated with clinical disease progression (19). As expected the nucleotide changes at position 5 of the Sp1III motif are different between subtype B (C5T) and subtype C (T5A). However, the effect of a single mutation at position five of the Sp1III motif in the HIV-1C LTR is not known.

We hypothesized that the Sp1 IIIT5A mutation may mediate HIV-1C LTR transcription activity, since this mutation occurs within the core-promoter, which is required to initiated viral gene transcription. Our data show the successful introduction of the single mutation (T5A) at position five of the Sp1III motif (Figure 1).

**Figure 1:**
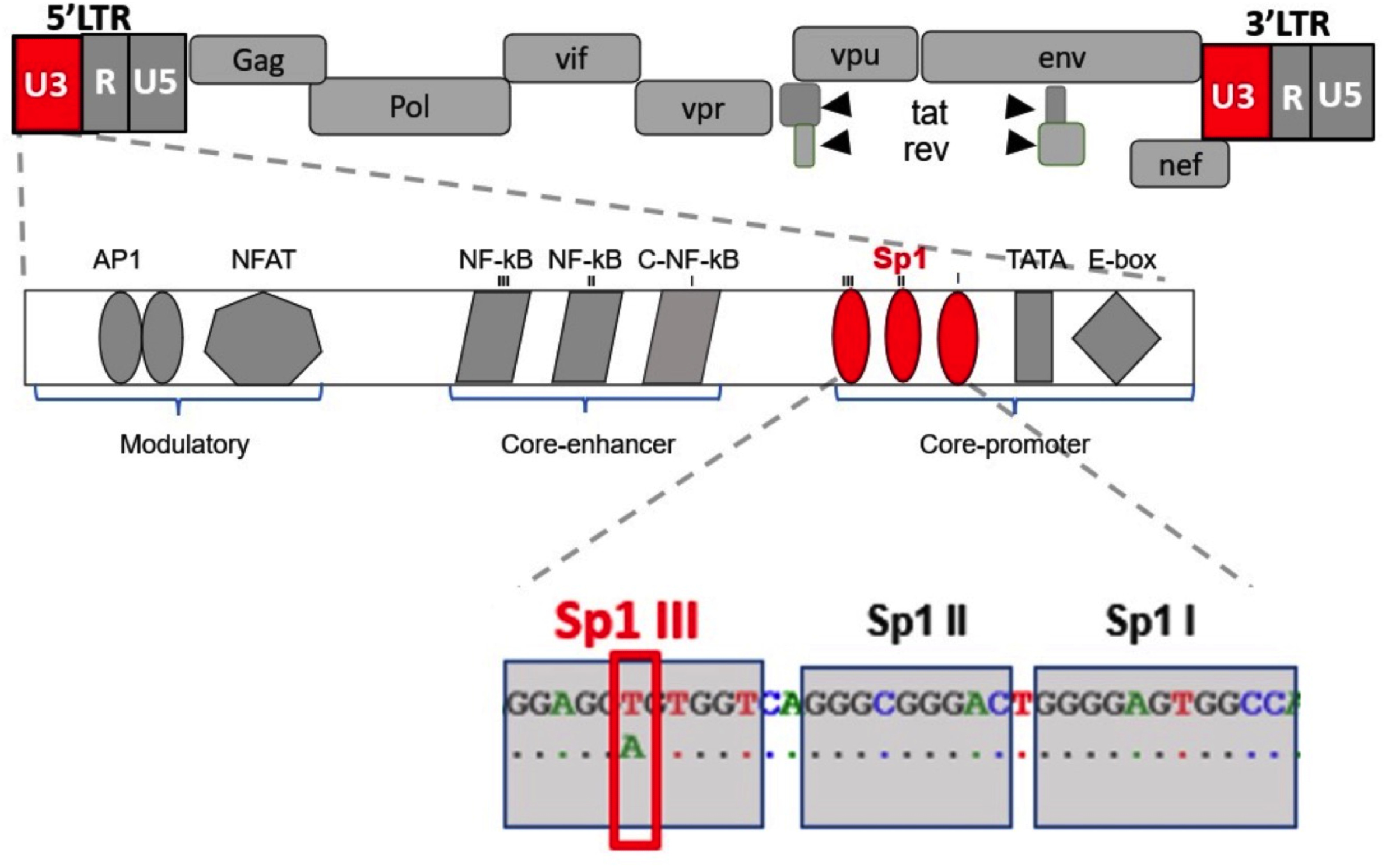
Sequence alignment of CSF-derived HIV-1C LTR with A mutation at position 5 of Sp1 III site (Sp1III5A). The LTR consists of modulatory, core-enhancer and core-promoter regions. The CSF-derived HIV-1C LTR with Sp1III5A was aligned with a Zambian subtype C reference (AF127567.1). The A mutation was successfully introduced at position 5 of the Sp1 III site located in the core-promoter region.

Specifically, we show that Sp1III5A mutant significantly increases basal transcription activity of HIV-1C LTR in both the SVG (p<0.0001) and Jurkat (p=0.0052) cells compared to the canonical Sp1III sequence, Sp1III5T (Figure 2A). Consistently, Sp1III5A mutant significantly enhanced Tat-mediated of HIV-1C LTR transcription activity in SVG (p<0.0001) as well as in Jurkat (p<0.0001) cell lines compared to HIV-1C LTR exhibiting the canonical sequence, Sp1III5T (Figure 2B). Notably, the Sp1III5A mutant had a more pronounced effect on basal and Tat-mediated HIV-1C LTR transcriptional activity in SVG compared to Jurkat cells (p<0.0001) (Figure 2A and B).

**Figure 2:**
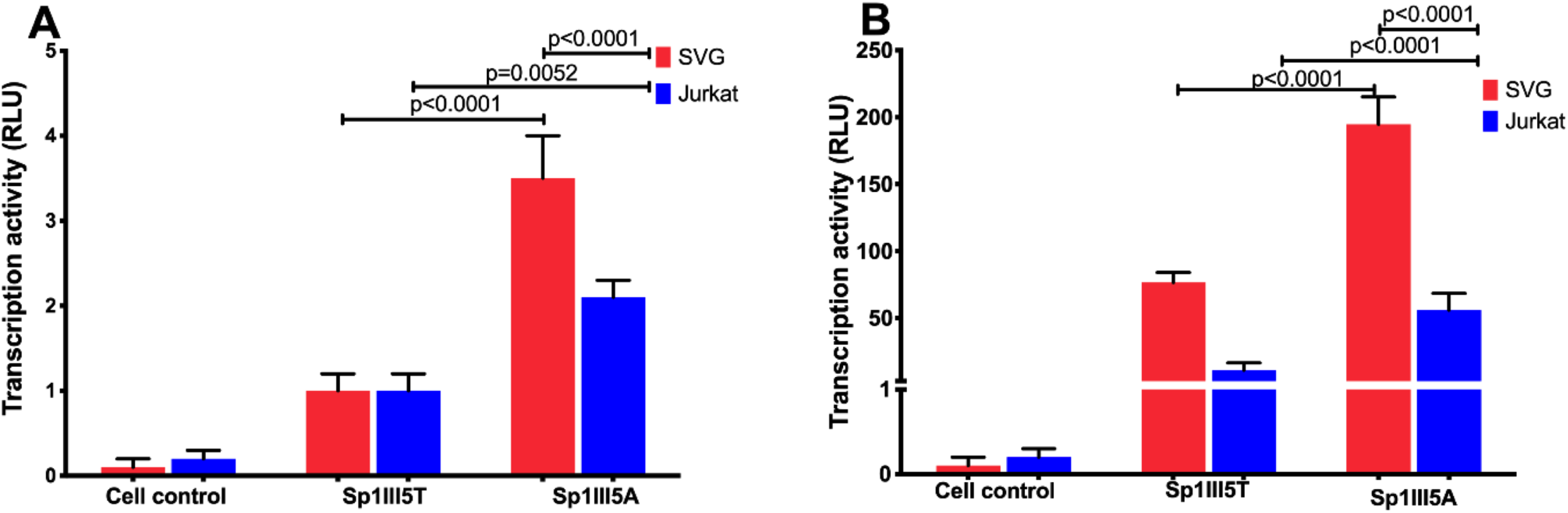
The transcription activity of HIV-1 subtype C LTR in Astrocytes and Jurkat cell lines. (A) Basal transcription activity of the HIV-1C LTR in SVGs (red) and Jurkat cell line (blue). (B) Tat-mediated transcription activity of the HIV-1C LTR in both SVGs (red) and Jurkat cell line (blue).

### Specificity protein 1 (Sp1) transcription factor expression levels of in SVG and Jurkat cell lines

According to earlier research, certain antiviral factors, like the *γ*-IFN-inducible protein 16 (IFI16) (23) and tripartite motif-containing protein 22 (TRIM22) (24), restrict the availability of cellular transcription factor Sp1, which is essential for effective viral gene expression, rather than targeting HIV-1 directly. Therefore, we hypothesized that significantly higher transcription activity observed in SVGs could be to these cells expressing higher levels of Sp1 transcription factor compared to Jurkat cells.

Our data show that the Sp1 expression levels were not significantly different between the SVG and Jurkat cells (p=0.0814), evident from their similar band intensity (Figure 3). However, we observed a trend that may correspond to the enhanced transcriptional activity in SVG cell line. Taken together these data suggest that Sp1III5A significantly increased LTR transcription activity observed in SVG and Jurkat cells may not be due to Sp1 expression levels but to other mechanism, such as its binding to the Sp1III motif.

**Figure 3:**
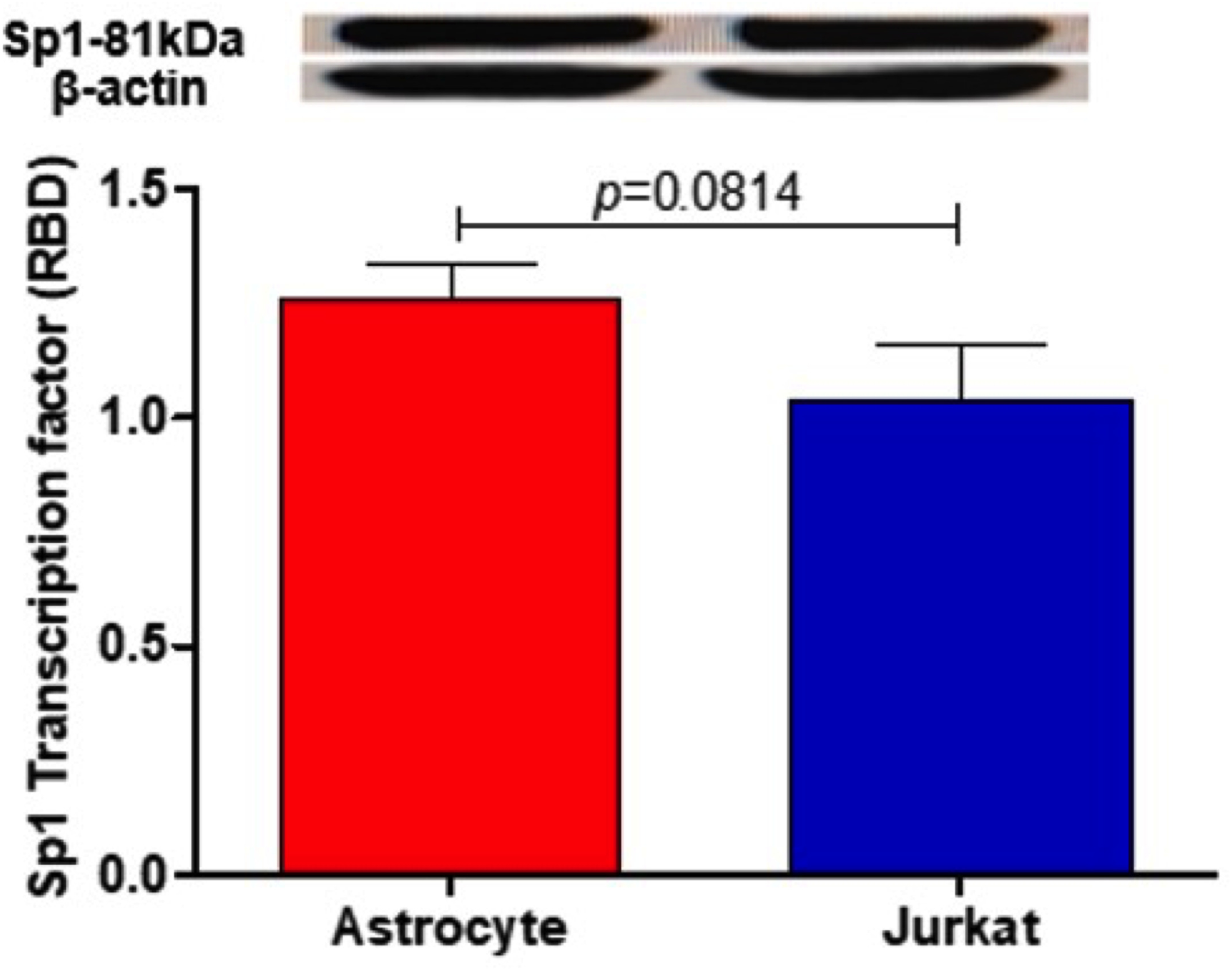
The expression levels of Sp1 transcription factor in SVG and Jurkat cell lines. Sp1 band intensity was similar across the two cell lines, SVG and Jurkat. B-actin was used as a house keeping gene. The red bar represent Sp1 expression levels in Astrocytes (SVG) while the blue bar represent Sp1 expression in Jurkat cells.

### Molecular Docking and Dynamics Simulation

#### Docking Calculations

The docking calculations aim to predict how a protein interacts with the nucleotides by simulating their binding modes and identifying the most likely conformations for complex formation. The results (Table 1) typically provide insights into the binding affinity, binding interface, and specific amino acid-nucleotide interactions.

**Table 1:**
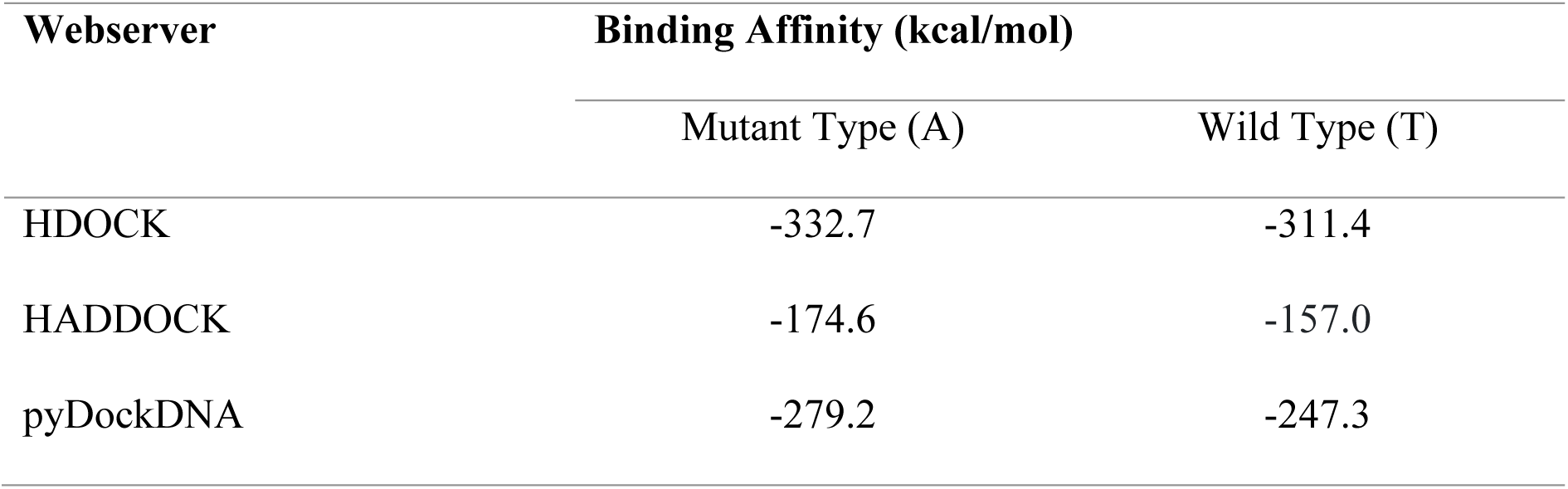
The binding affinity of the wild and Mutant types of the LTR against the Specificity protein 1.

#### Molecular Dynamic Simulation

Molecular interactions, conformational flexibility, and environmental factors interact to control protein-DNA stability during molecular dynamics (MD) simulations. The integrity of the complex is maintained by important stabilizing interactions such as van der Waals forces, hydrophobic contacts, electrostatic interactions, and hydrogen bonds. These connections can be strengthened or weakened by dynamic conformational changes in both DNA and proteins. Temperature and solvent content are two other environmental variables that affect stability. To effectively describe these dynamics, the simulation duration and force field selection are crucial. The Sp1III5A mutant showed stronger stability and protein-DNA interactions as revealed by root mean square deviation (RMSD). Specifically, the quantification of the stability of the Sp1 protein interaction with different LTRs in molecular dynamics (MD) simulations RMSD yielded RMSD values of 17.96259 Å for the LTR Sp1IIIT and 13.02608 Å for the LTR Sp1IIIA suggesting that the stability of the two complexes and structural dynamics differ (Figure 4). The docking pose of the three web servers was examined. An alignment analysis was performed which showed a very low deviation (less than 0.7 Å (RMSD)) (Figure S3).

**Figure 4:**
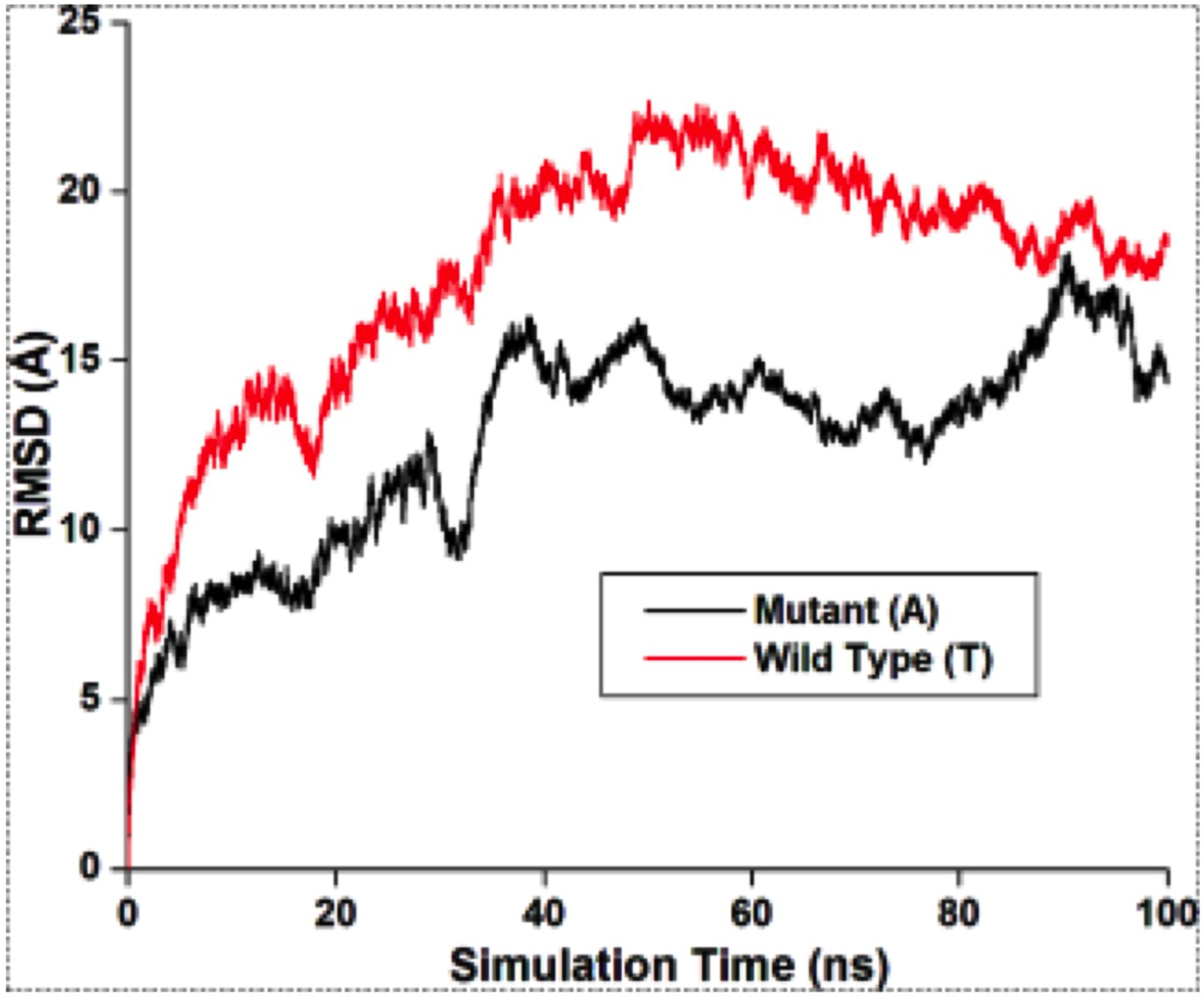
The Root Mean Square Deviation (RMSD) plot of the SP1 in complex with LTR Mutant and Wild type.

#### Molecular Interaction

Protein-DNA molecular interactions are essential for the regulation of gene expression, DNA replication, repair, and overall genomic stability. These interactions can be sequence-specific, where proteins such as transcription factors recognize specific DNA base pairs, or non-specific, where proteins interact primarily with the DNA backbone. Key forces, including hydrogen bonds, electrostatic forces, and van der Waals interactions, help stabilize these complexes. Our data highlights the active residue of Sp1 protein interacting both the Sp1IIIT (canonical Sp1III sequence) and Sp1IIIA mutant The table (Table 2).

**Table 2:** Interaction of the active site residue of SP1 with the wild type and Mutant.

| Protein Residues | Wild type (T) | Mutant(A) |
| --- | --- | --- |
| LYS153 | Charge-Charge | N/A |
| HIE156 | Conventional hydrogen bond | N/A |
| ARG168 | Salt-bridge, Charge-charge | Salt-bridge, Charge-charge |
| LYS179 | Charge-charge | Charge-charge |
| THR182 | Carbon hydrogen bond and<br>Conventional hydrogen bond | N/A |
| ARG183 | Conventional hydrogen bond | Conventional hydrogen bond<br>and Pi-Cation |
| ASP185 | Conventional hydrogen bond | N/A |
| GLU186 | N/A | Carbon hydrogen bond |
| ARG189 | N/A | Conventional hydrogen bond |
| LYS207 | LYS207 | Salt-bridge, Charge-charge |
| ARG211 | N/A | Conventional hydrogen bond |
| HIE214 | Conventional hydrogen bond | Carbon hydrogen bond and Pi-Pi Stacked |
| LYS217 | N/A | Pi-Alkyl |
| ARG275 | Charge-charge | N/A |
| GLY298 | Carbon hydrogen bond, | N/A |

The structural interactions between the LTR Sp1III5A and canonical Sp1III5T DNA-SP1 complex show the overall structure of each complex, highlighting changes in amino acid interactions with DNA (Figure 5). Specifically, the regions of the DNA where the interactions take place in Sp1III position five where T to A mutation occurs. In the mutant structure, an adenine base forms a notable interaction with Asp185, while in the canonical Sp1III5T sequence (wild-type) structure, a thymine base interacts with Arg189. These structural differences likely indicate changes in binding specificity or stability caused by the mutation, which may affect the biological function of the DNA-protein complex. Therefore, high binding affinity of the mutant type is because of collective interaction of residue of the LTR (mutant).

**Figure 5:**
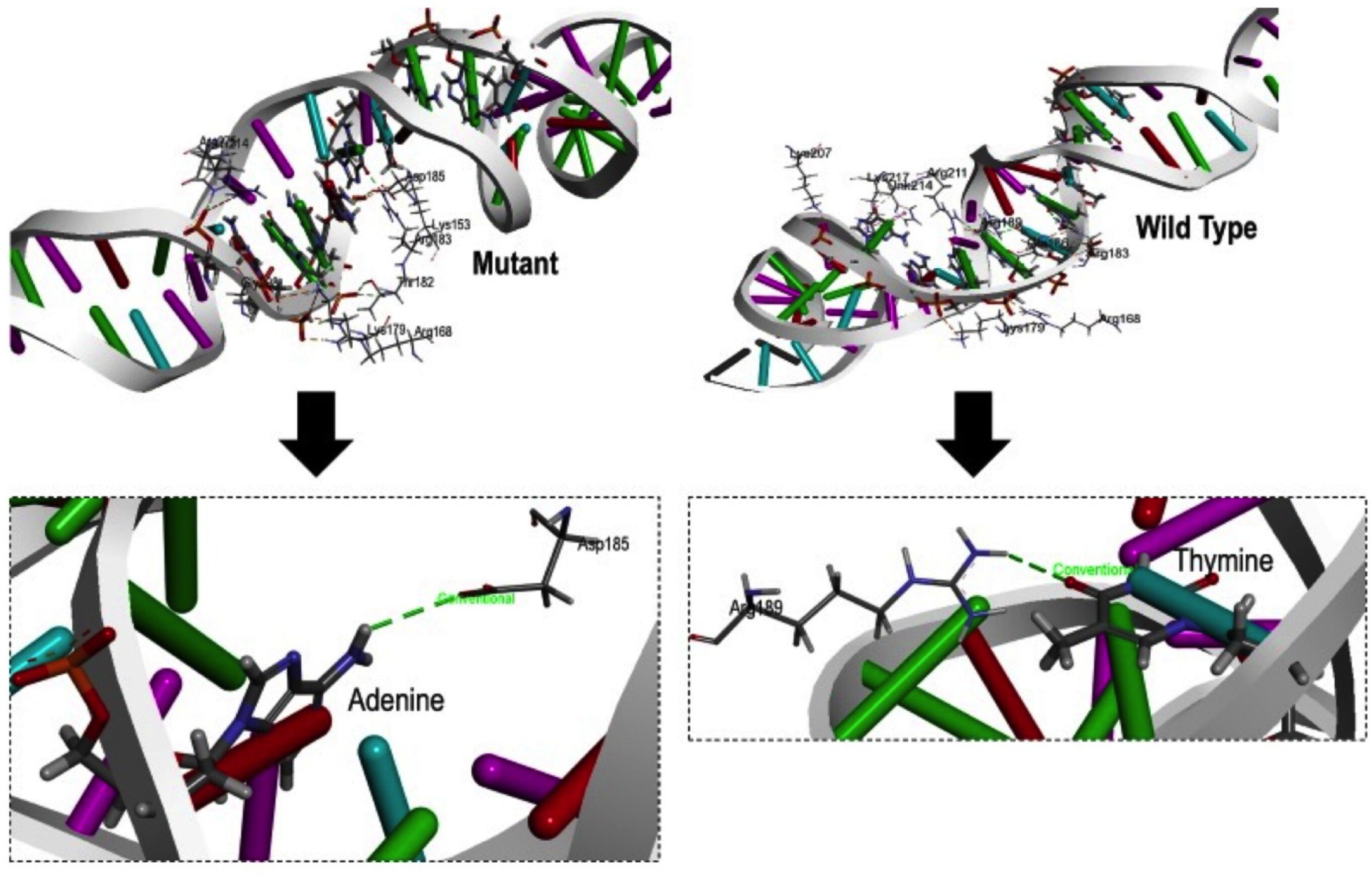
Interaction of LTR (mutant and wild type) with SP1 protein, displaying interaction of the mutant residue (Thymine to Adenine) at the binding site of LTR.

## DISCUSSION

The genetic variation of HIV-1C subtype T/F viruses in the LTR regions, particularly the higher frequency of single nucleotide polymorphisms (SNPs) in the Sp1III motif, significantly impacts transcriptional activity and disease outcome (22). While previous studies have highlighted this correlation, the individual contributions of specific mutations within this motif remain unclear. In this study, we hypothesized that the Sp1IIIT5A mutation in the core-promoter region of HIV-1C LTR independently mediates the LTR transcription activity and Sp1 transcription factor binding affinity. Our findings are consistent with this hypothesis, demonstrating that Thymine (T)-to-Adenine (A) substitution at position five of Sp1 III binding site (Sp1IIIA5T) significantly increases both basal and Tat-mediated HIV-1C LTR transcription activity in SVG and Jurkat cell. Notably, this enhancement was more pronounced in SVG cells, despite similar levels of Sp1 expression across both cell types.

These data are consistent with earlier studies linking genetic variation within the Sp1 motifs to dysregulated LTR transcription activity and disease outcome (18, 22, 25, 26). For example, earlier study reported that peripheral blood mononuclear cells-derived HIV-1 LTRs from individuals with rapid disease progression frequently contained a fourth Sp1 motif, while those from long-term non-progressor exhibited variable Sp1III motifs with diminished Tat-mediated transcription activity (25). Similarly, mutations at other Sp1 and C/EBP motifs, such as Sp1III5T and C/EBP3T, have been shown to differentially regulate LTR-driven gene expression, with their combination resulting in reduced transcription activity (18). Furthermore, a comparative study of HIV-1B’ variants circulating in China revealed that a deletion at position five in the Sp1III motif enhanced transactional activity compared to HIV-1B circulating in America, which retained a Guanine at this position (26). Importantly, while these prior studies largely focused on subtype B, our study addresses HIV-1C, the most globally prevalent subtype, responsible for approximately 50% of infections worldwide and 98% in southern Africa (21).

Interestingly, intracellular defence proteins, known as host restriction factors, such as Tripartite motif 22 (TRIM22) are capable of limiting the amount of Sp1 levels and inhibiting HIV-1 replication (24) but in this study the Sp1 levels were the same in different across different cell lines. This suggests that a different mechanism may be involved in reduced transcription activity by LTR Sp1III5A.

Our findings contribute to this discourse by showing that the Sp1IIIT5A mutation in HIV-1C enhances Sp1 binding affinity, as evidenced by molecular docking analyses. Compared to the canonical Sp1III5T sequence, Sp1III5A displayed stronger binding, likely due to more stable protein-DNA interaction. Computational docking tools (HDOCK, HADDICK, and pyDockDNA) consistently indicated enhanced binding affinity for the Sp1IIIT5A mutation, despite algorithms (27–29). This stronger interaction was characterized by reduced conformational flexibility (lower RMSD values), suggesting greater stability of the mutant Sp1III5A DNA-Sp1 complex. Further molecular analyses highlighted the role of charge-charge interactions and hydrogen bonding in stabilizing the Sp1IIIt5A-Sp1 complex. Key residues such as LYS153, LYS179, and ARG168 established strong electrostatic attractions, while hydrogen bonds involving HIE156, ARG183, and ASP185 reinformed structural integrity. Notably, ARG183 exhibited a dual role in the Sp1IIIT5A complex through hydrogen bonding and pi-cation interactions, contributing to enhanced specificity and stability compared to the canonical sequence. Additional interactions, including stacking and hydrogen bonding, involving residues such as THR182 and ARG189, further supported the robustness of the mutant complex.

## CONCLUSION

In this study, we show that the Sp1IIIT5A mutation alone significantly enhances HIV-1C LTR transcriptional activity by increasing Sp1 binding affinity and complex stability. These findings provide new insights into the molecular mechanisms underpinning HIV-1C transcriptional regulation and highlight the potential role of Sp1 motif variations in influencing disease progression. Future research should investigate the effect of Sp1IIIT5A mutation on latent reservoir establishment and reactivation. Understanding how the Sp1IIIT5A mutation interacts with host cellular factors involved in latency could inform potential therapeutic strategies that target these regulatory pathways to mitigate HIV-1 pathogenesis and address the challenges posed by latent reservoirs. Understanding transcriptional regulation and latency mechanisms, will inform future studies on the development of cure strategies aimed at achieving virus clearance or long-term control, remission.

## Materials and Methods

### Ethics statement

The study was approved by Biomedical Research Committee (BREC) of the University of KwaZulu Natal (BREC/00006386/2023).

### Construction of recombinant HIV-1C consensus and Sp1IIIT5A LTR U3R-pGL3 luciferase basic reporter vectors

The HIV-1C consensus LTR sequence (amplified from the following reagent obtained through the NIH HIV Reagent Program, Division of AIDS, NIAID, NIH: Human Immunodeficiency Virus-1 (HIV-1) 93ZM74 LTR Luciferase Reporter Vector, ARP-4789, contributed by Dr. Reink Jeeninga and Dr. Ben Berkhout) or Sp1IIIT5A LTR U3R designed and purchased as gene blocks (Whitehead Scientific, South Africa, Cape Town) were cloned into the pGL3 Basic vector expressing Firefly Luciferase (Promega, Corporation, Wisconsin, USA) as we previously described (22). Briefly, pGL3 Basic vector and consensus or Sp1IIIT5A LTR U3R PCR product were digest independently with *Kpn*I and *Hind*III (New England Biolabs, Ipswich, MA, USA) according to the manufacturer’s instructions. The restriction fragments were analyzed on a 1% agarose gel to confirm the restriction fragment sizes. The correct size fragments corresponding to the consensus LTR U3R region and the linearized pGL3 Basic vector were purified by gel extraction and purified using GeneJet Gel Extraction Kit (ThermoFisher Scientific, Waltham, MA, USA) as per the manufacturer’s instructions. Digested consensus or Sp1IIIT5A LTR U3R region was ligated into the linearized pGL3 Basic Vector/plasmid using 1U of T4 DNA Ligase (New England Biolabs, MA, USA), following the manufacturer’s instructions, to create the recombinant consensus or Sp1IIIT5A LTR U3R-pGL3 Luciferase reporter plasmid. The recombinant plasmids were transformed into JM109 competent *E*. *coli* cells (Promega, USA, Madison) according to the manufacturer’s instructions. The transformed cells were plated on ampicillin agar plates and incubated overnight at 37°C for 14 hours. A randomly picked colony was used to inoculate the culture, and plasmids were purified using the GeneJet Plasmid Mini Prep Kit (Invitrogen, Carlsbad, CA) as per the manufacturer’s instructions. The presence of the correct insert (consensus or Sp1IIIT5A LTR U3R region) in the recombinant plasmid was confirmed by sequencing using the Big Dye Terminator v3.1 Cycle Sequencing Reaction (Applied Biosystems, California, USA). Sequencing samples were analyzed using the ABI 3130xl Genetic Analyzer. Large-scale stocks of the recombinant consensus or Sp1IIIT5A U3R-pGL3 Luciferase reporter plasmids were generated using the Plasmid Maxi Kit (Qiagen, Valencia, CA) for immediate use or long-term storage at -80°C.

### Analysis of reporter gene expression

Astrocyte (SVG) and T-lymphocyte (Jurkat) and cell lines were transiently transfected using Polyethylenimine (PEI) transfection reagent (ThermoFisher Scientific, Waltham, MA, USA), as described previously by our group and others (22, 30). Briefly, SVG cells were seeded into 96-well flat-bottom tissue culture plates at a density of 5,000 cells/well in 100 μL while Jurkat cells were seeded in a 24-well tissue culture plates at a density of 5 x 10^5^ cells/well in 400 μL of antibiotic free media supplemented with 10% fetal bovine serum (FBS). The following day, cells were transfected with a DNA pool of 400 ng, consisting of 300 ng of the recombinant consensus or Sp1IIIT5A LTR-pGL3 plasmid and 100 ng of recombinant control insert DNA-pTarget plasmid (Promega, Madison, WI) was prepared for Tat-minus transfection. Alternatively, for Tat-transfected conditions, 100 ng of recombinant HIV-1C consensus plasmid (pcDNA3.1(+) expressing isogenic mutant HIV-1 subtype C BL43/02 Tat, ARP-11785, contributed by Dr. Udaykumar Ranga) cloned into pTarget plasmid was prepared in 50 μL of serum-free media. In a separate tube, 1 μL of PEI (Polyplus, NY) was mixed with 49 μL of serum-free RPMI medium to prepare the lipid transfection reagent. The transfection complex, 50 μL of the PEI-RPMI mixture was mixed with 50 μL of the recombinant plasmid pool containing consensus or Sp1IIIT5A LTR-pGL3 plasmid and recombinant control or Tat-pTarget plasmids. The 100 μL plasmid-lipid mixture was incubated at room temperature for 20 minutes and the transfection mixture was added to the corresponding wells of the plates containing either SVG or Jurkat cells. Twelve hours post-transfection, the cells were washed to remove lipid complexes and resuspended in the corresponding volume (100 μL for SVG and 500 μL for Jurkat) of R10 medium. The transfected cells were incubated for an additional 24 hours prior to the Luciferase assay. Luciferase expression was measured using the Bright-Glo Luciferase Assay System (Promega, Madison, WI). Briefly, 150 μL of the cell culture was mixed with 100 μL of the Bright-Glo substrate reagent in a black round-bottom 96-well plate. The luciferase activity was recorded using the Victor Nivo Multimode plate reader (PerkinElmer, Massachusetts, USA). The experiments were conducted in triplicate wells, and each experiment was repeated at least twice.

### Western blotting

Protein expression of Sp1 in Jurkat and SVGs was determined using Western blotting assay. Briefly, Jurkat and SVG cell culture was independently lysed in a cocktail of protease and phosphatase inhibitors supplemented with Cytobuster^TM^ protein extraction Reagent (Sigma-Aldrich, St. Louis, Missouri, USA). The cells were stored on ice for 30 minutes before being scraped mechanically. Cellular lysates were centrifuged (10000 x g, 10 minutes, 4°C, using Eppendorf, Hamburg, Germany) and the supernatants with the crude protein extract were aspirated into fresh 1.5 mL micro-centrifuge tubes and kept on ice.

The crude protein was quantified using the bicinchoninic acid (BCA) assay. The astrocyte and Jurkat proteins were then standardized to concentrations of 1.166 mg/mL and 0.854 mg/mL, respectively and the standardized samples were produced in Laemmli buffer [dH_2_O, 0.5M Tris– HCl (pH 6.8), glycerol, 10% SDS, 5% beta-mercaptoethanol, and 1% bromophenol blue] by boiling for 5minutes at 100°C. Using the Bio-Rad compact power supply, the protein samples and the molecular weight marker were electrophoresed (1h, 150V) in sodium dodecyl sulphate polyacrylamide gels (10% resolving gel and 4% stacking gel) and transferred to a nitrocellulose membrane using the Bio-Rad Transblot^®^ Turbo^TM^ Transfer System (Bio-Rad, Hercules, California, USA). The membranes were blocked for 1 hour (RT) with gentle shaking in 1:5 000 (5% BSA/non-fat dry milk (NFDM) in Tris-buffered saline with 0.05% Tween 20 (TTBS; 150mM NaCl, 3mM KCl, 25mM Tris). The membranes were probed overnight with primary antibody (anti-Sp1); 1:1,000 in 5% BSA/TTBS (Cell Signalling Technology, Danvers, Massachusetts, USA) at 4°C.

Membranes were rinsed five times in TTBS for five times (10 minutes each) before being incubated with the secondary antibody for 1 hour. The Clarity Western ECL Substrate Kit (Bio-Rad, Hercules, California, USA) was used to visualize protein bands and were detected using the Molecular Imager® ChemiDoc^TM^ XRS Bio-Red Imaging System (Hercules, California, USA). The membranes were quenched with 5% H_2_O_2_ (30 minutes, 37°C), washed three times with TTBS (10 minutes, RT), blocked in 5% BSA in TTBS (1h, RT), and re-probed with HRP-conjugated secondary antibody for the house-keeping protein, β-actin (Sigma Aldrich, St. Louis, Missouri, USA) after detection (1:5,000 dilutions in 5% BSA, 30 minutes, RT). Images were captured and analyzed by dividing the band intensity of each sample by the respective loading controls to determine relative band density (RBD).

### Modelling and Validation of the LTR and Protein (Specificity protein 1)

Owing to the absence of a resolved crystal structure for the LTR (nucleotide) and the Sp1, the nucleotide was modeled using two different webservers: model.it^®^ Server (http://pongor.itk.ppke.hu/dna/model_it.html#/modelit_form) (31) and Supercomputing Facility for Bioinformatics & Computational Biology webserver (http://www.scfbio-iitd.res.in/software/drugdesign/bdna.jsp#). The web servers are used to model nucleotide sequences (wild type and mutant) that integrate computational biology and structural modeling. This is done by inputting a nucleotide sequence of the DNA in the correct format (A, T, C, G) and then selecting parameters like DNA type and structural conformation. The server generates a 3D model, often allowing for energy minimization and refinement to ensure stability. The structure obtained from both servers shows a similar orientation (visually and sequentially) (shown in Figure S2 in the supplementary material). The protein structure was modeled using the amino acids sequence generated from the GenBank database (accession code: ACR22508) (https://www.ncbi.nlm.nih.gov/genbank/) (32). The accession code corresponded to an Alphafold-predicted protein structure on the UniProt database (33) and AlphaFold Protein Structure Database (https://alphafold.ebi.ac.uk/) (34, 35) with the access code C4PGM0 shown in Figure S1A (cyan). The Protein was further modeled (homology modeling) using the SWISS-MODEL (https://swissmodel.expasy.org/) (36). The homology model of Specificity protein 1 demonstrated an excellent sequence coverage of 100% and sequence similarity of 96.66%, with the highest sequence identity being 94.95%. The model quality was affirmed by GMQE and QMEANDisCo global model evaluation scores of 0.51 and 0.39, respectively, suggesting a highly reliable structure. The predicted 3D structure showed predominant regions of high and a few regions of low confidence, indicated by blue and red colors, respectively (Figure S1B). Additionally, the QMEAN Z-score of -9.28, which is below to zero, further indicated the stability and accuracy of the model (Figure S1C). The spatial distribution of the modeled structure was consistent with that of a non-redundant set of PDB structures, reflecting a good local similarity to the target (Figures S1D and S1E). The Ramachandran plot analysis further supported the model high quality, showing a MolProbity score of 1.65, a clash score of 0.23, and 82.83% of residues in the favored regions with 6.06% outliers. Moreover, there were only 3.02% rotamer outliers, 9 C-Beta deviations, 4 bad bonds out of 2227, and 56 bad angles out of 3009 (Figure 1F) (37). These findings collectively underscore the robustness and reliability of the homology model for Specificity protein 1. A sequence alignment analysis was performed on the two structures (the AlphaFold predicted structure (C4PGM0) and the homology modeled structure) using CLUSTAL O (1.2.4) multiple sequence alignment (38) which indicates that both structures have similar sequence orientation (Figure S2 Supplementary material). The two structures were analyzed visually using Chimera (39) and Discovery Studio (40) and it was observed that the homology result has a well-oriented secondary structure compared to the AlphaFold prediction. The homology model was selected for further analysis.

### Molecular Docking Calculation

The docking calculations were done using three different web servers namely; HDOCK (http://hdock.phys.hust.edu.cn/) (41), HADDOCK (https://rascar.science.uu.nl/haddock2.4/) (42), and pyDockDNA (https://model3dbio.csic.es/pydockdna) (28) webservers. HDOCK is particularly useful for protein-DNA docking due to its efficient combination of template-based modeling and free docking, allowing it to explore a wide range of potential binding poses. This versatility can help identify multiple viable interactions, which is important when dealing with complex biomolecular systems. HADDOCK, on the other hand, emphasizes the incorporation of experimental data into the docking process. If you have any structural or biochemical data about the protein-DNA interaction, HADDOCK can utilize that information to refine the docking results. This data-driven approach often leads to more accurate and reliable models, particularly in scenarios where you have specific binding site information. pyDockDNA is tailored specifically for DNA and RNA interactions, focusing on the unique characteristics of nucleic acids. Its algorithms take into account the distinct binding geometries and electrostatic properties involved in DNA-protein interactions, which can enhance the accuracy of the docking predictions. Using these three web servers, gives insight for comparison and validate your docking results, ensuring good complexity of the protein-DNA (Sp1-LTR) interactions. The structure of the protein and the LTR (both mutant and wild type) were supplied to the webservers in PDB format and the results were visualized using Chimera and Discovery studio

### Molecular Dynamics Simulation

The Amber 18 PMEMD engine GPU version was used to run a 100 ns molecular dynamics (MD) simulation on Sp1 complexes of the LTR wild and mutant forms (43). The Amber18 Leap module neutralized the complexes, and an orthorhombic TIP3P water box was used for simulation. Using the conjugate gradient algorithm, energy minimization in 10,000 iterations with a 500 kcal/mol^2^ constraint (44). The complexes were heated progressively over 10 ps from 0 to 300 K using static atoms and volumes to determine the heating process. The Berendsen barostat mechanism, which had a harmonic restriction of 10 kcal/mol·Å² and a collision frequency of 1.0 ps, kept the system pressure at 1 bar. The operating temperature (300° K) was maintained during the 10 ns, 5,000,000 steps systems equilibration. An isobaric isothermal ensemble (NPT) was maintained by maintaining a constant number of atoms and pressure (45). To keep the pressure at 1 bar, the Berendsen barostat was used. Hydrogen atom bonds were created using the SHAKE algorithm. To examine stability, the AMBER18 GPU PTRAJ module computes the root mean square deviation (RMSD) after analyzing the coordinates and trajectories every 1 ps (46).

## Acknowledgments

Research reported in this publication was supported by the South African Medical Research Council with funds received from the National Department of Health (MRC-RFA-SHIP 02-2018 to PM), a grant from the National Research Foundation Thuthuka Funding Instrument (TTK160529166617 to PM) and Poliomyelitis Research Foundation (Grant number: 23/66 to PM and grant number: 23/40 to NM). This study was funded by the Poliomyelitis Research Foundation (Grant number: 23/40). The work was also supported in part by grants from the Bill and Melinda Gates Foundation [OPP1212883 and INV-033558]; Gilead Sciences, Inc [Grant ID #00406] and the International AIDS Vaccine Initiative (IAVI) [UKZNRSA1001 to TN]. This work also received funding through the Sub-Saharan African Network for TB/HIV Research Excellence (SANTHE) [grant # DEL-15-006]. SANTHE receives additional funds from the Science for Africa Foundation with support from Wellcome Trust and the UK Foreign, Commonwealth & Development Office and is part of the EDCPT2 programme supported by the European Union; the Bill & Melinda Gates Foundation [INV-033558]; and Gilead Sciences Inc., [19275]. All content contained within is that of the authors and does not necessarily reflect positions or policies of any SANTHE funder. For the purpose of open access, the author has applied a CC BY public copyright licence to any Author Accepted Manuscript version arising from this submission. Lastly, the study was also partly funded by Erasmus+ which is the EU’s programme to support education, training, youth and sport in Europe [Erasmus+KA107].

---

**Figure S1:**
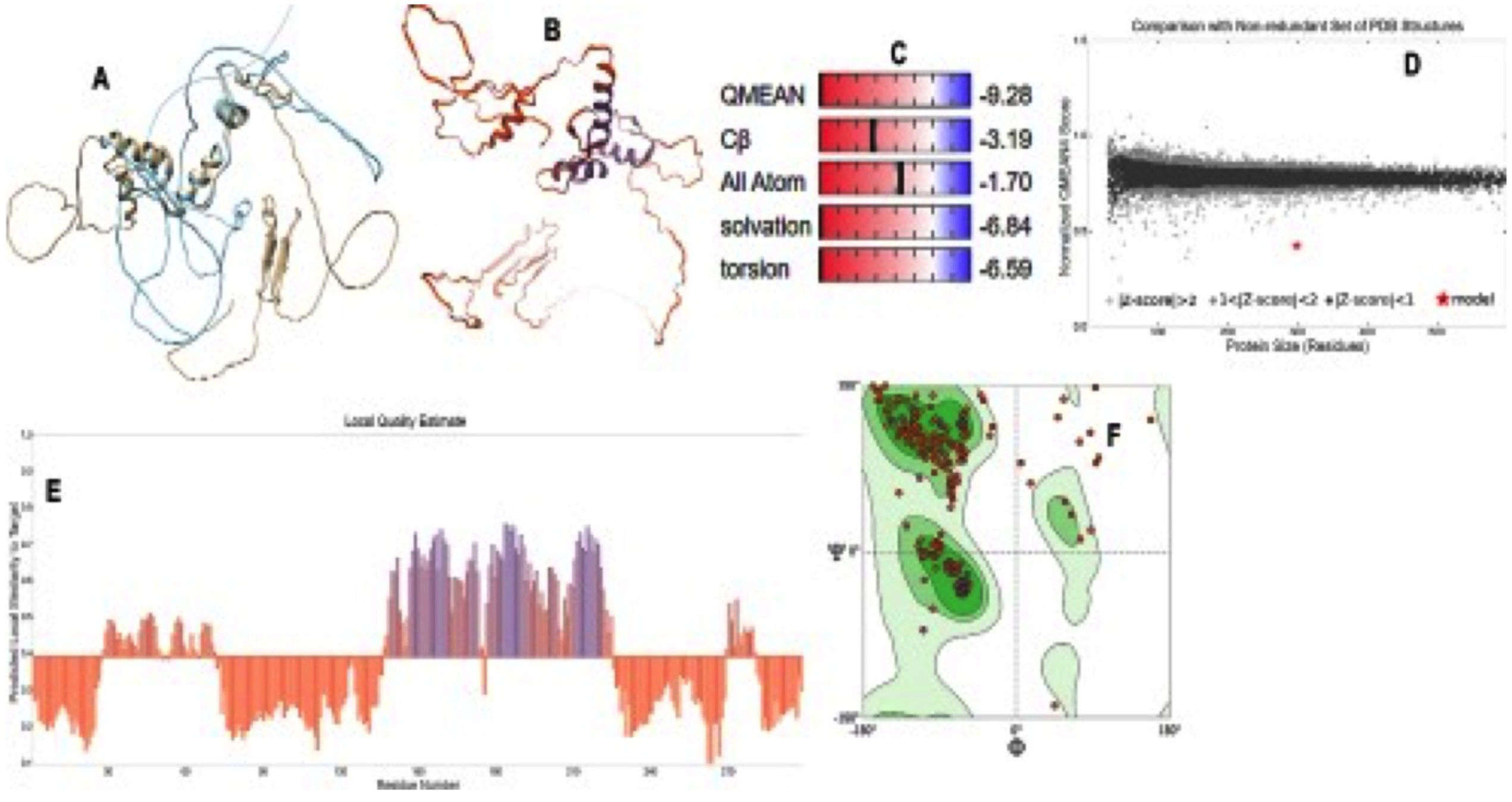
Validation of homology model from SWISS-MODEL web server showing the alignment analysis of C4PGM0 and model structure (A), The computed three-dimensional (3D) homology model colored by the confidence gradient scale Alignment analysis of C4PGM0 and model structure (B), Global quality estimate scores (C), Graph of Comparison with a non-redundant set of PDB structures (D), Local quality estimate graph (E), and The Ramachandran plot analysis of the modeled structure (F).

**Figure S2:**
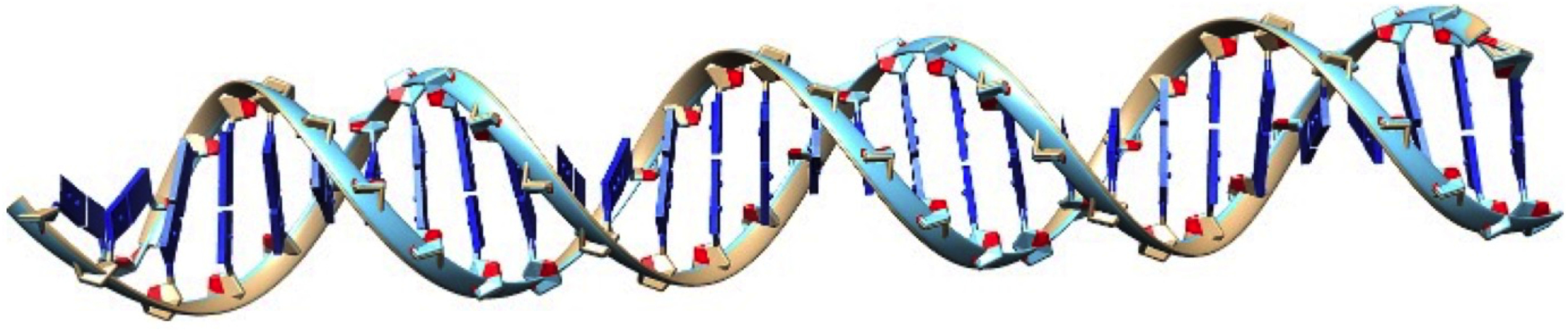
Alignment of the two modeled LTR nucleotide structure.

**Figure S3:**
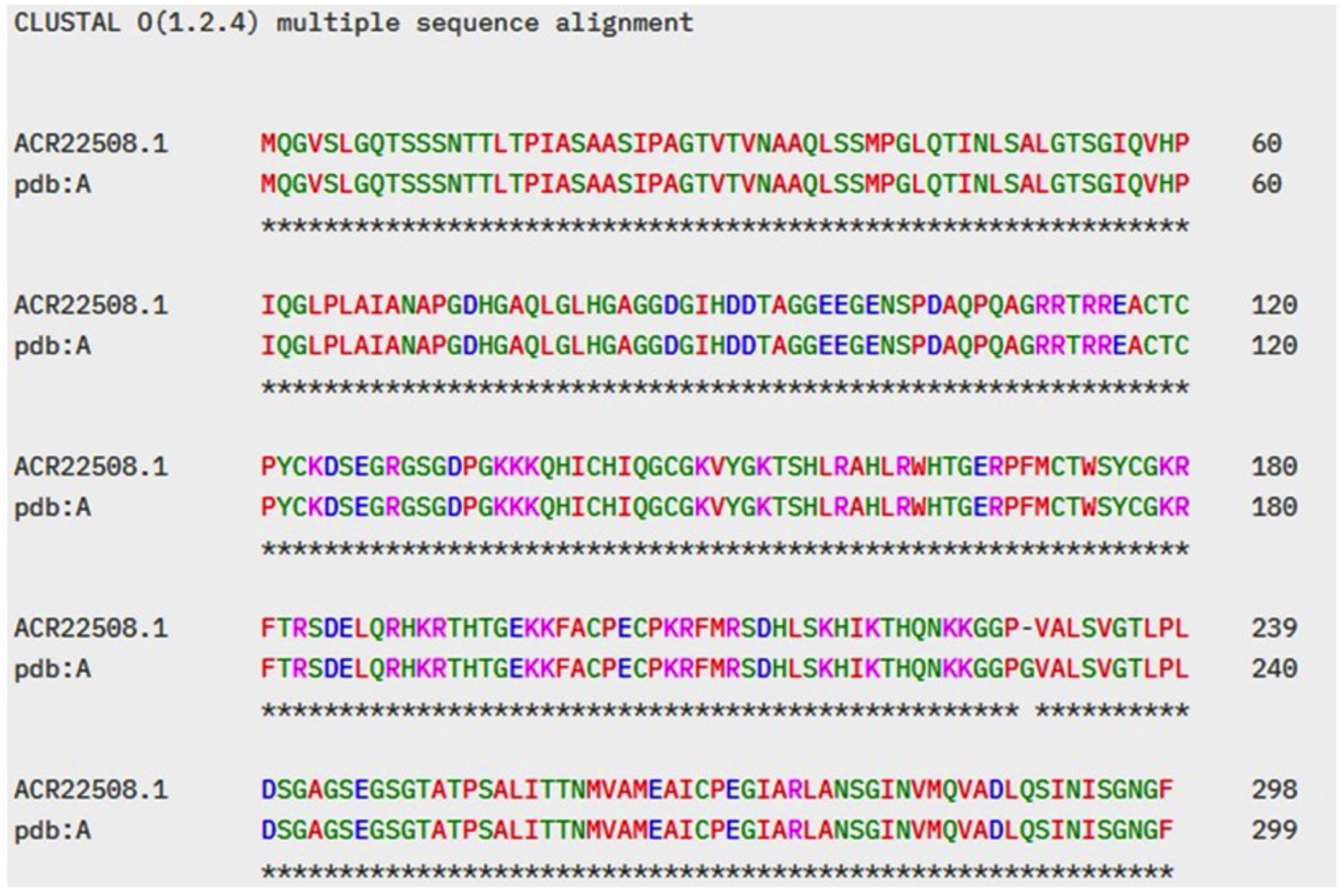
CLUSTAL O (1.2.4) multiple sequence alignment of the AlphaFold predicted structure (ACR22508.1) and the homology modeled structure.

**Figure S4:**
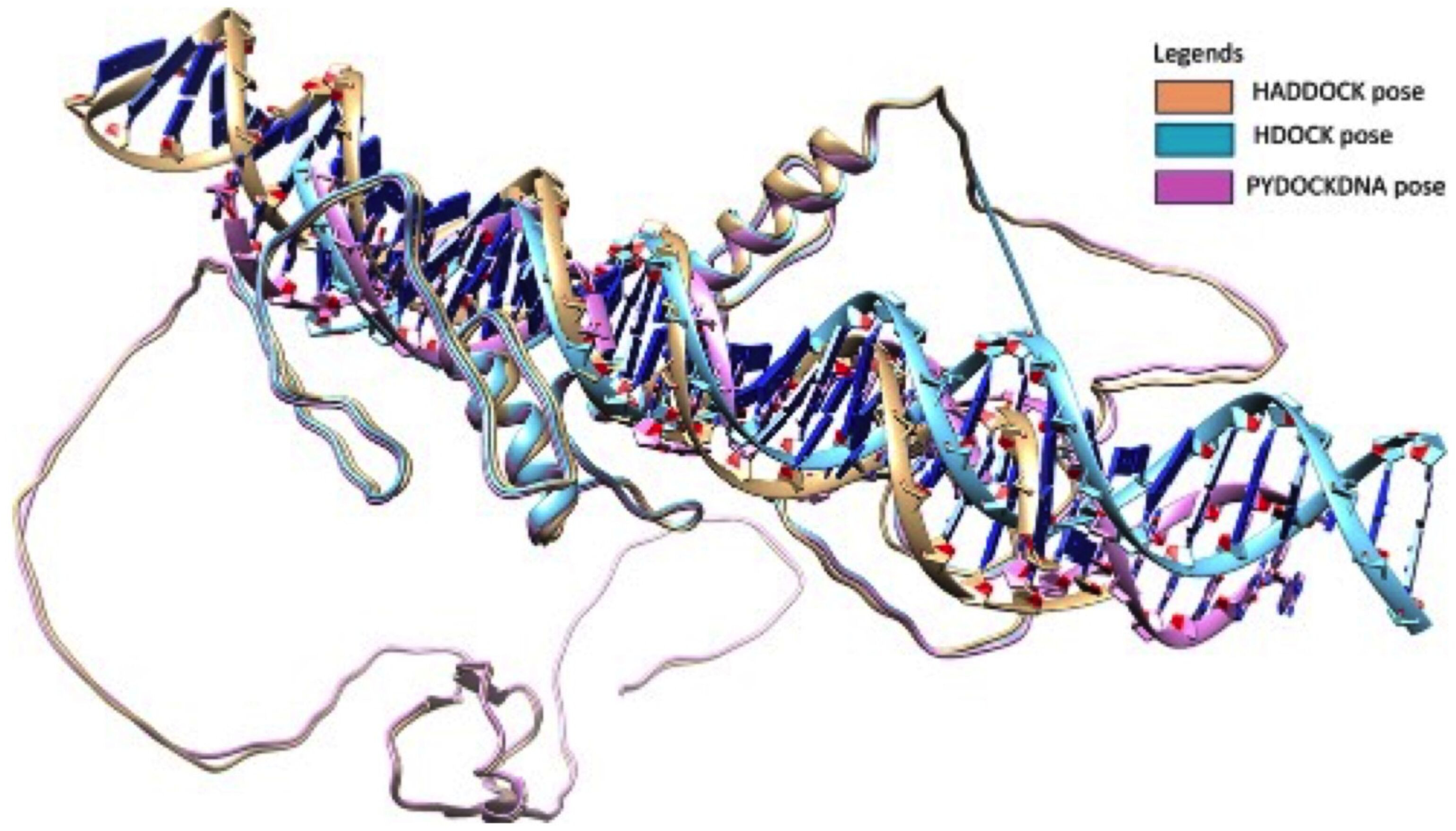
Alignment of the docking pose of the best pose of the Sp1-LTR complexes from three web servers.

